# 5-HAYED peptide can protect AD brain by scavenging the redundant iron ions and the catalyzed radicals

**DOI:** 10.1101/373639

**Authors:** Zhenyou Zou, Shengxi Shao, Li Li, Xiaoying Cheng, Qiqiong Shen

## Abstract

Alzheimer’s disease (AD) is a progressive neurodegenerative disease characterized by memory and cognitive decline. It is incurable currently and <u>places a great burden on the caregivers</u> of patients. Iron is rich in the brain of AD suffers. It catalyzes radicals which impairs neurons. Therefore, reducing the redundant brain iron is pressing to ease AD. To scavenge the excessive brain iron catalyzed radical, thus protect the brain and decrease the incidence of AD. We synthesized a soluble iron-pro 5-HAYED peptide. By injecting 5-HAYED to the cerebrospinal fluid (CSF) of the AD mouse, we observed that the 5-HAYED is able to decrease the brain iron and radical level, which behaving neurons protection, and can ameliorate the cognition status for AD mouse. Further, 5-HAYED can decreased the AD incidence and can reverse the AD associated anemia and inflammation without hurt kidney and liver.

## 1. Introduction

Alzheimer’s disease (AD) is a progressive neurodegenerative disease characterized by memory and cognitive decline [1]. It is incurable currently and places a great burden on the caregivers of patients [2].

The cause of AD is not well understood. Many articles have demonstrated that increased oxidative stress in AD brains may have a pathogenic role in neuronal degeneration; they impair membrane lipids and proteins and ultimately result in neuronal death [3, 4].

Iron is a transitional metal. It can catalyze free oxidative radical generation by deoxidizing oxygen to oxyradicals or by donating electron to hydrogen peroxide, forming hydroxyl radicals [5]. Our previous study revealed that AD brains have higher levels of iron than the normal brains, and the iron distribution determines the regional density of radicals [6]. Therefore, reducing redundant iron in the brain may be an option to ease AD.

Metal-chelators, such as desferrioxamine (DFO) and deferiprone (DFP), have been used as AD treatments in clinical trials [7]. These agents have slowed the progression of AD in certain cases [8, 9]; however, the fundamental aspects of their biochemistry have severely limited their effectiveness. For example, the hexadentate iron chelator DFO can cause fever, hearing loss and severe allergic reactions [10]; the lipid-soluble iron chelator DFP can lower neutrophils and white blood cell count which causing life-threatening infections [11, 12]. Therefore, designing or identifying a nontoxic iron chelator that removes excessive iron from the brain is pressing.

Amyloid beta (Aβ) peptide consists of 36–43 amino acids and is cleaved by beta secretase from amyloid precursor protein (APP) [13]. This peptide possesses high affinity Cu^2+^ and Fe^3+^ binding sites at Asp (D) _1_, Glu(E)_3_, and His(H)_6,13,14_ [14]. By binding metal ions, Aβ aggregates to form amyloid plaques in AD brains [15] and reduces Cu^2+^ to Cu^1+^ and Fe^3+^ to Fe^2+^, catalyzing the O_2_-dependent production of H_2_O_2_ [16]. Based on the iron-binding characteristic of Aβ, we combined the metal-pro amino acids histidine (H), glutamic acid (E) and aspartic acid (D) to an oligomer. In addition, to enhance the solubility of the peptide, we added the polar amino acid tyrosine (Y) and the smallest nonpolar amino acid alanine (A). And at last, we synthesized a 5-repeat HAYED dissoluble oligomer.

To examine the iron chelating and radical purging efficiency, we administrated it to the iron-rich medium cultured SH-SY5Y neuroblastoma cells and then treated it to the wild aged Kunming (KM) mice, which display symptoms similar to those human patients with AD, such as memory deterioration, neurofibrillary tangles, gliocyte hyperplasia and inflammation, and in particular, the brain iron level in these mice was higher than that of the normal level [17], which is highly suitable for examining the brain radical level and damage and performing the cognitive assays.

## 2. Results

### AD mouse has higher level of iron and hydroxyl radicals and observed neuron necrosis in brain

AD model mouse brain has more iron than that of the normal. To be specific, AD samples have 0.68 μg iron per gram of brain tissue in average, whereas, the normal samples have only 0.32 μg per gram brain (Fig. 1-A). The topography (Fig. 1-A) reveals that iron is richer in the corpus columella, cortex, hippocampus and cingulate cortex of AD brain, but relatively scarce in the amygdala and ventricles. By contrast, in most areas of the normal brain, except for sporadic abundance in the ventricles, columella, cortex and CA1, iron is relatively lower; the hippocampus is the rarest zone.

Accordingly, hydroxyl radical levels in the AD brain were 70% higher than those of the normal (Fig. 1-B). Neuron necrosis is widespread, indicating by the Tunnel stained brown area in Fig. 1-C.

**Fig. 1.**
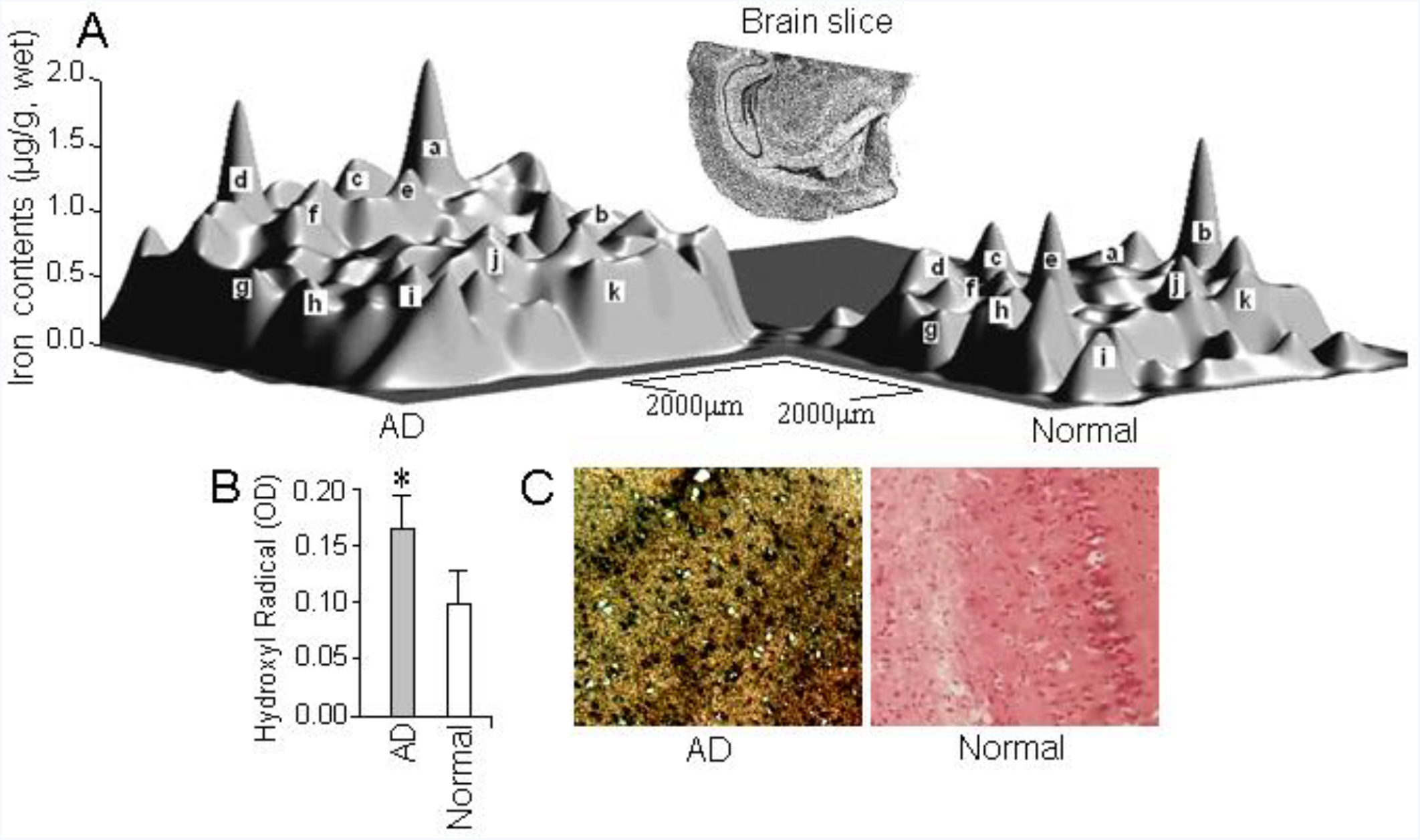
AD brain has more iron and hydroxyl radicals and damaged. **A)** Iron distribution in the brain. The AD brain has more iron than the normal brain. *a)* IG-RSGb; *b)* D3V; *c)* PtA; *d)* S1BF; *e)* CA1; *f)* CA2-CA3; *g)* Temporal cortex; *h)* PRh-LEnt; *i)* Amygdala; *j)* Thalamus; *k)* Hypothalamus; **B)** Overall, hydroxyl radical levels are high in the AD brain; **C)** The TUNEL assay shows dark staining in the AD brain, indicating a necrosis.

### 5-HAYED can bind iron

A mass spectrometer (Fig. 2-A) shows that the particles with 5 hydrions appear at the peak 715.85, 4 hydrions particles and 3 hydrion participles locate at 894 and 1192 respectively. With the regularity, the molecular weight of the whole synthesized oligomer is 3574.5, which is appropriate for the 5-HAYED repeats peptide. Using a protein sequencer (PPSQ-21A/23A, Shimatsu Corp., Japan), the peptide oligomer was sequenced as “HAYED HAYED HAYED HAYED HAYED”, which is approximately 300 nm in length (Fig. 2-B-c) and completely conformed to our design.

**Fig. 2.**
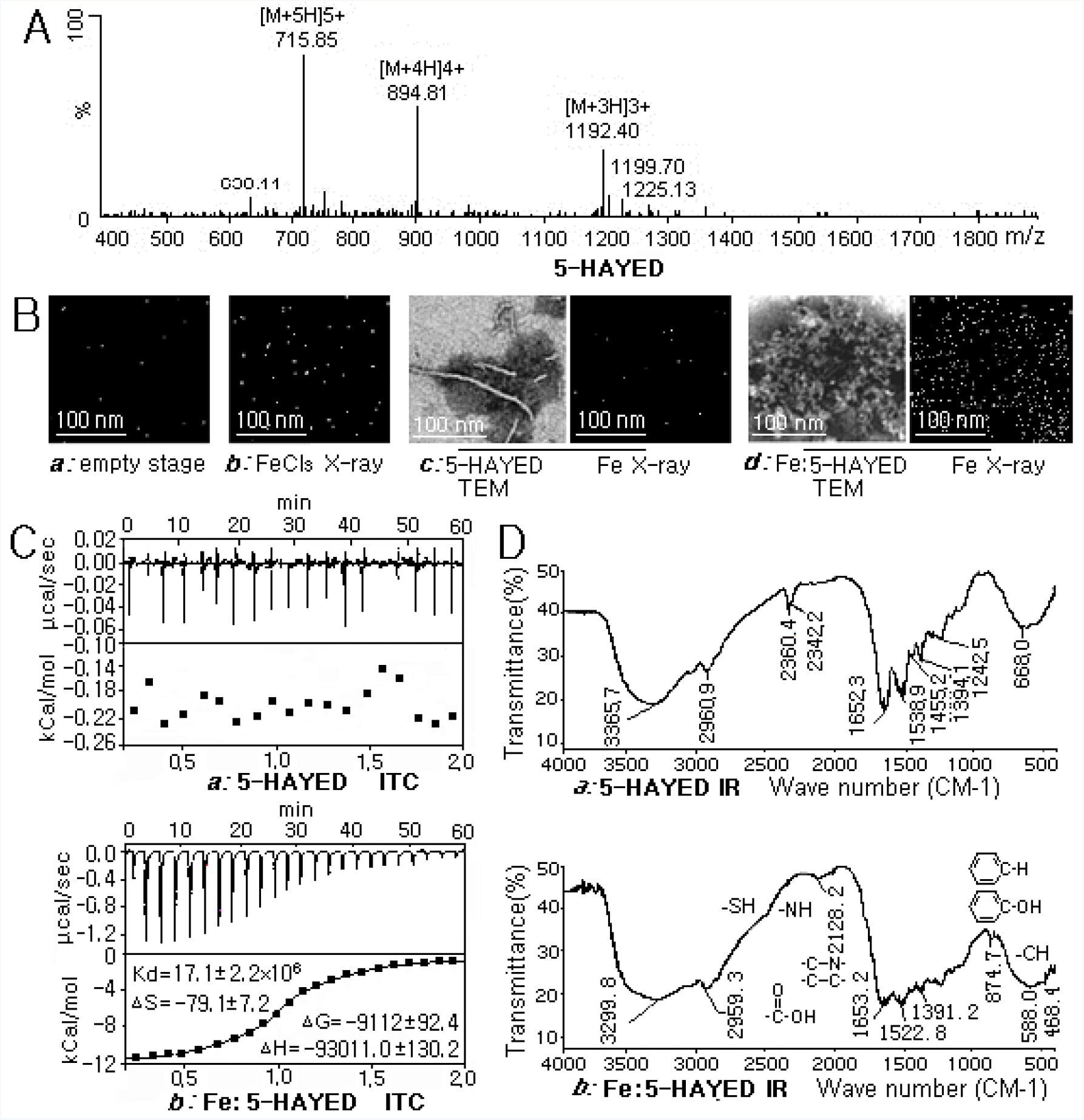
5-HAYED is affine to iron ions. **A)** Mass spectrum of the synthesized 5-HAYED amino acid oligomer. The site m/z=715.85 indicates the particles with 5 hydrions; 894 is the peak of 4 hydrion particles, and 1192 is the site of 3 hydrion particles. The molecular weight of the entire peptide is 3574.5. **B)** TEM image of 5-HAYED oligomers with or without FeCl_3_ incubation. The FeCl_3_ incubation resulted in the agglomeration of the 5-HAYED oligomer, and X-ray energy spectrum shows iron iso-directed along the amino acid fibril. **C)** Isothermal titration calorimetry shows that the enthalpy of the Fe: 5-HAYED compound lost 93011 kCal/mole enthalpy and 9112.6 kCal/mole Gibbs, indicating that 5-HAYED has a higher affinity for iron. **D)** Infrared spectra of the 5-HAYED before and after the FeCl_3_ incubation. The signals of phenyl-OH (720 cm^-1^) in the Tyr residue,-COOH (2925 cm^-1^ and 1600~1300 cm^-1^) in the Glu and Asp residues, and C-N (2375 cm^-1^) and C=N (700~1615 cm^-1^) in the His residue were weakened, suggesting that iron binding occurred. Furthermore, the signals representing the backbone acylamide N-H stretching (3500~3100 cm^-1^), the C=O bond shift (1680~1630 cm^-1^), the N-H bending (1655~1590 cm^-1^) and C-N stretching (1420~1400 cm^-1^) were transformed, indicating that a tortuosity occurred in the backbone after the iron incubation.

Pure 5-HAYED oligomers display liner-like fibrils (Fig. 2-B-c). But after incubated with FeCl3, the oligomers were observed curled and conglomerated (Fig. 2-B-d), and the iron was detected iso-directed along the 5-HAYED fibrils (the bright spots in the right panel of Fig. 2-B-d representing the Fe positions), which strongly suggesting the junctions exist between the iron atoms and the amino acid oligomers.

Isothermal titration calorimetry revealed that 5-HAYED is high affinity to iron. Shown by Fig. 2-C-b, after the FeCl_3_ titration, the 5-HAYED lost 93011 kCal/mole in enthalpy and 9112.6 kCal/mole in Gibbs energy. No binding occurs to 5-HAYED if only titrated with ITC buffer (Fig. 2-C-a). Specifically, 5-HAYED oligomer bind irons at the residues His, Tyr, Asp, and Glu, for the peaks in the band of 1600~1300 cm^-1^ in infrared chromatography, which representing the C=O group vibrations in the carboxyl of Glu and Asp, changed after the 5-HAYED incubated with the FeCl_3_; and the peak 720 cm^-1^, which representing the weakening of the phenyl hydroxide of Tyr (Fig. 2-D-b); the absorption apex at 2375 cm^-1^, which representing the stretching of the C-N bond, and the peaks in the 1700~1615 cm^-1^ band, which displaying the stretching of the C=N bond in the imidazole ring of the histidine residue, were all weakened (Fig. 2-D-b), committed by the chelating of iron atoms.

### 5-HAYED can protect cells by reducing iron catalyzed radicals

The SH-SY5Y cells, if incubated in iron-rich medium (containing 0.016 M FeCl_3_·6H_2_O, imitating AD CSF) for 36 hours, they atrophied, with the axons and dendrites shrunk, and dendritic spines decreased (Fig. 3-A-a). However, if 100 pM 5-HAYED was pre-added in the medium, the cells survived with axons and dendrites extending and spines rich on dendrites (Fig. 3-A-b). Furthermore, the cells exhibit active cytoplasmic transport, for transport cysts can be seen along the dendrites (pointed by the arrow in Fig. 3-A-b). By contrast, in the iron-stressed cells, no transporting cysts can be found. It is obviously that the iron-stress damaged the cell physiological activities.

**Fig. 3.**
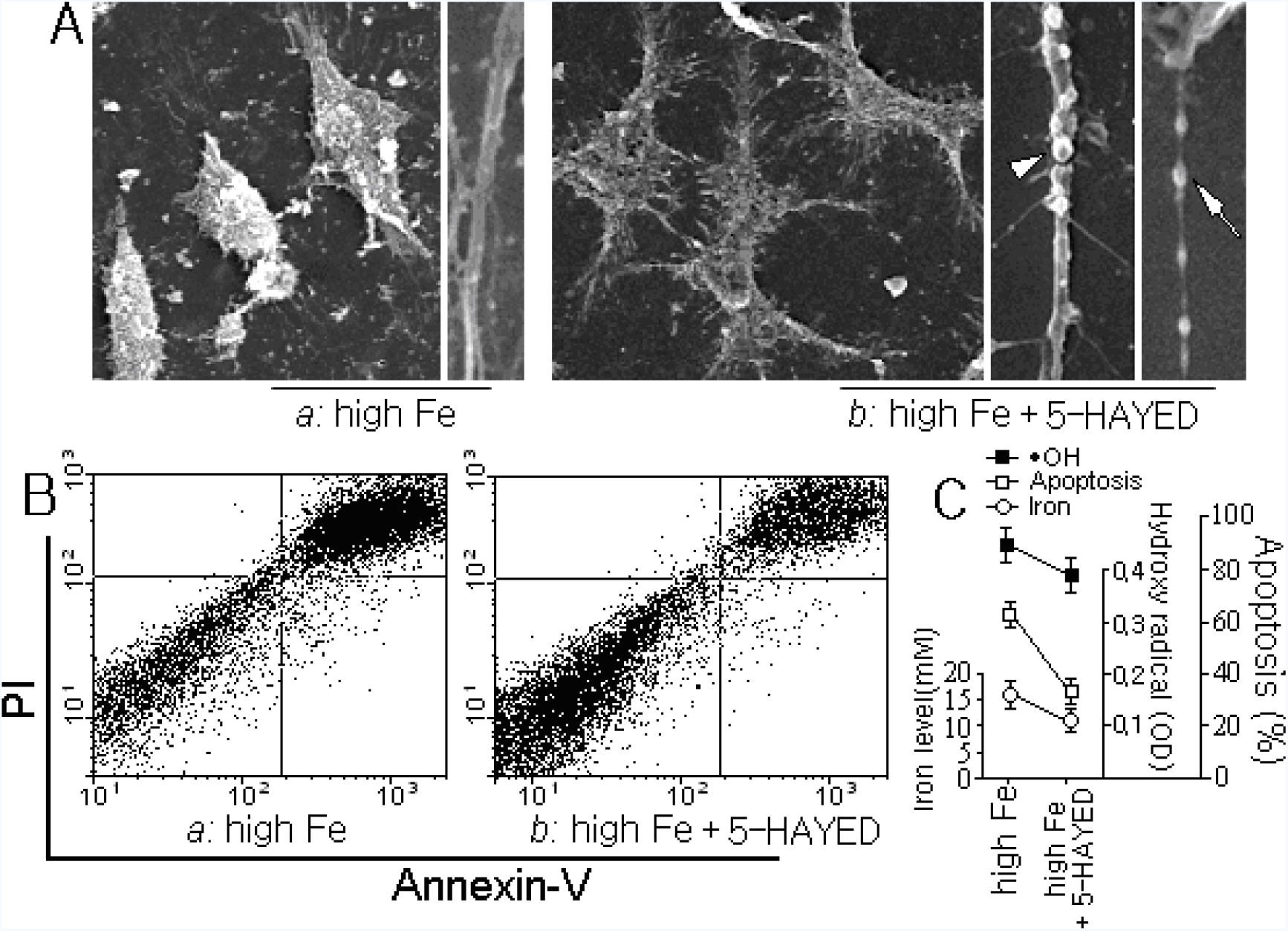
5-HAYED can reduce redundant iron ions and hydroxyl radical thus protects cultured cells. **A)**SH-SY5Y cells shrunk in the iron-rich medium, the dendritic spines (Δ) decreased, and axonal transportation (↖) disappeared; By contrast, in 5-HAYED supplemented high iron medium, the cells survived, the axons and dendrites extended, the dendritic spines increased, and the cytoplasmic transport was active. Transport cysts were observed along the dendrites. **B and C)** 5-HAYED peptide reduced hydroxyl radicals and free iron in the medium, resulting in increased cell survival.

Spectrophotometry and ICP assays revealed that 5-HAYED peptide has the ability to clear hydroxyl radicals and to chelate the redundant iron (Fig. 3-C). The hydroxyl radicals decreased from 0.886 to 0.766, and free iron ions in the medium decreased by one third. As a result, more cells survived in the medium, shown by Fig. 3-B.

### 5-HAYED behaves neuron protection and can ameliorate cognitive statue for AD suffers

The wild aged KM mice, which were used as the Alzheimer’s disease model, many neurons in the cortex and hippocampus could be observed injured by Tunnel assay (indicating by the green spots in Fig. 4-A). The iron and hydroxy radical in the brain were 17 mM and 0.14 OD respectively, more than 70% and 180% higher than that of the normal (Fig. 4-B). However, when 5-HAYED was injected into the CSF, the free iron and the hydroxy radical decreased to 12 mM and 0.07 OD respectively (Fig.4-B), 30% and 1 times lower than that of the AD ones. And after the 5-HAYED treatment, the nerves in the AD brain appeared more organized and less necrosis (Fig. 4-A). It is obviously that 5-HAYED taken effect in neuron protection by reducing iron catalyzing radicals.

**Fig. 4.**
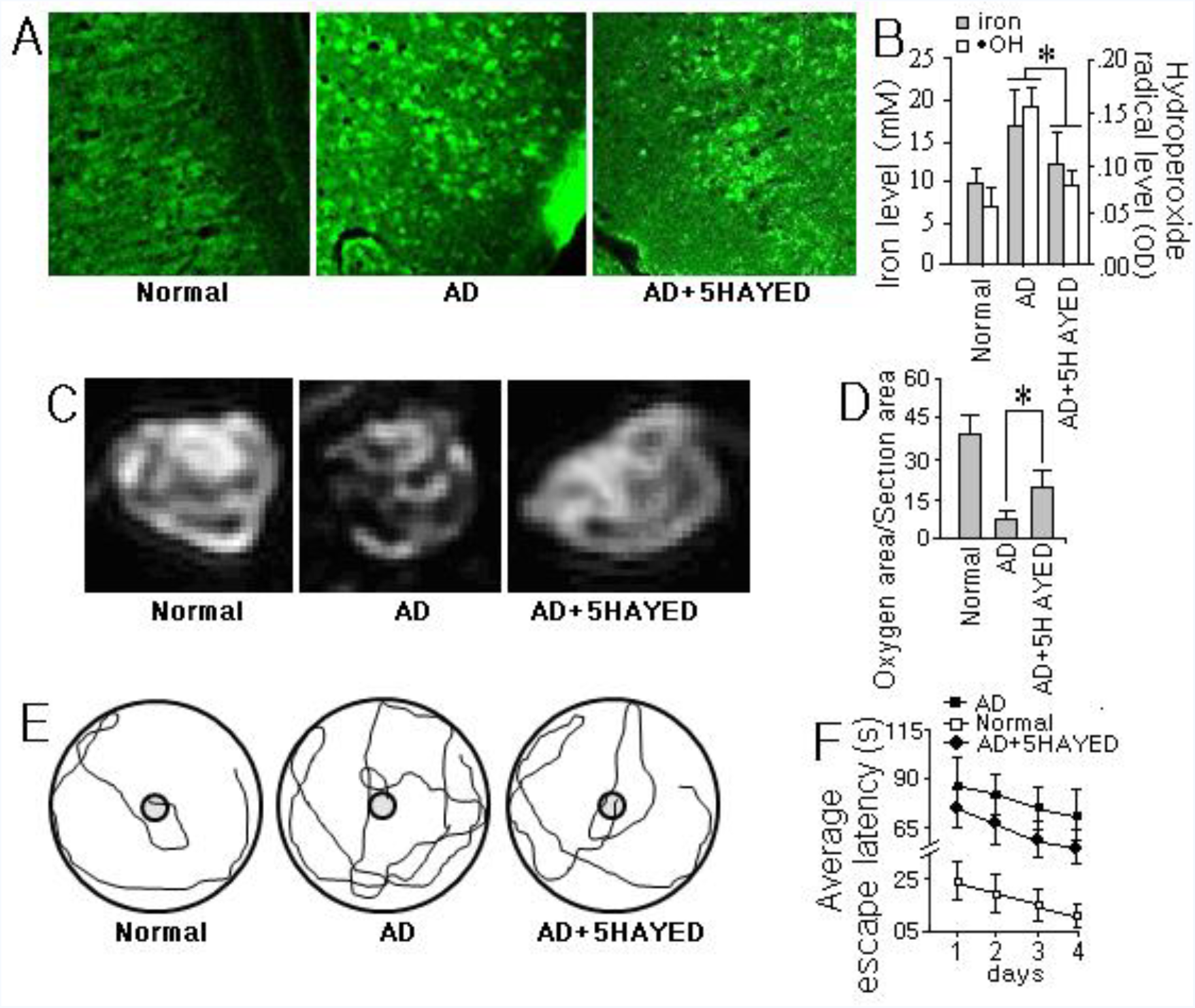
5-HAYED behaves neuron protection and cognitive statue amelioration properties and can lower AD incidence. **A)** TUNNEL assay shown that the AD brains treated with 5-HAYED display fewer necrotic neurocytes (the bright green spots) than the untreated AD brains, indicating that 5-HAYED has the function protecting brain. **B)** The iron and hydroxyl radical levels in CSFs. 5-HAYED treated brain has lower iron and hydroxyl radical level than that in the untreated AD group. **C** and **D)** fMRI shows that the area of blood oxygen metabolism in the 5-HAYED-treated brain occupied approximately 23% of the whole section, which is 1.5 times wider than that of the untreated aged wild AD mice and approximately restored half of the normal mice in area. **E** and **F)** Morris water maze assay shows that the distance that the 5-HAYED-treated AD mice swum to find the platform is obviously shorter than that swum by the untreated AD individual. The time that the 5-HAYED treated AD mice spent about 59 s in average to swim back to the underwater stage, 23 s saved compared with that of the untreated AD ones. *N=6; *: p <0.05.*

Even more, after the 5-HAYED treated, we found that the blood oxygen metabolism level increased by 152% compared with the untreated AD suffers (bright areas in the brain shown of Fig. 4-C and D). The Morris water maze assay further revealed that after 4 days of learning, the 1 month 5-HAYED treated mice behave less clumsy than the untreated wild aged ones. They spend 59 s on average to return to the platform, nearly 23 s saved than that of the untreated ones (Fig. 4-E and F), that is, the synthesized 5-HAYED obviously ameliorated the cognition status for the AD suffers.

### 5-HAYED transgenic mouse has low AD incidence

When 5-HAYED encoding oligonucleotide sequence was combined to the mouse genome (Fig. 5-B show the southern blot of 5-HAYED DNA) and expressed the 5-HAYED peptide (the brown spots in Fig. 5-C), AD rarely attacks the mouse (Fig. 5-D).

**Fig. 5.**
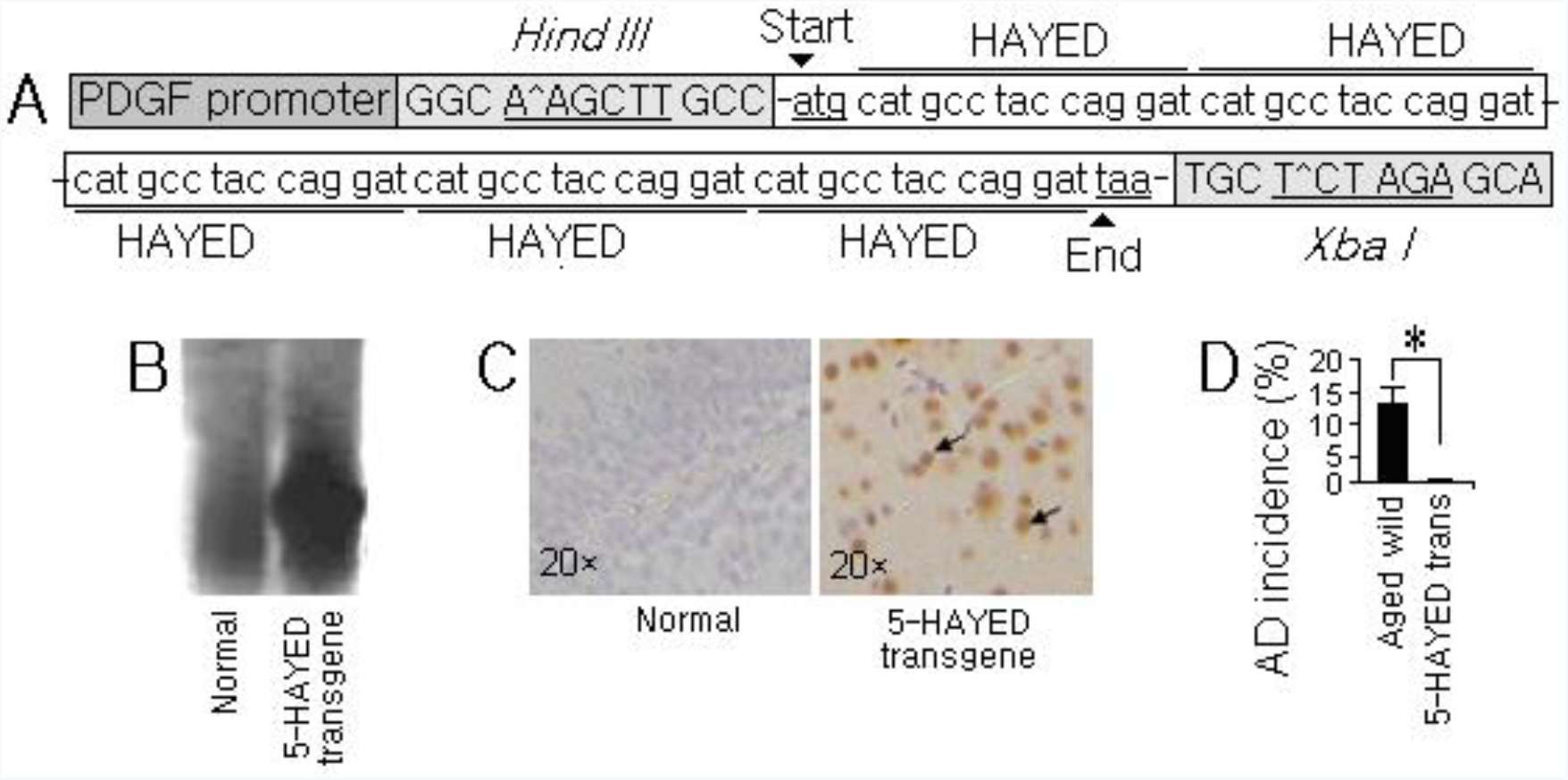
5-HAYED transgenic mouse has a lower AD incidence. **A)** Gene structure of 5-HAYED. PDGF promoter was combined upstream to initiate the transcription; restriction sequences of Hind III and Xba I were inserted for recombination; **B)** Southern blot of 5-HAYED in the DNA extracted from the transgenic mouse tail and **C)** immunohistochemistry of 5-HAYED peptide in the brain section of 5-HAYED transgenic mouse show the transgenic mouse expressed HAYED. **D)** 5-HAYED transgenic mouse has low AD incidence compared with the wild aged mice. *N=30; *: p <0.05.*

### 5-HAYED has no observed side effect on liver, kidney and blood

A good medicament should not only be high effective but should also be non-toxic to the human body. The clinic chemistry and blood test demonstrate that no obvious side effects attacked to the liver, kidney and blood of the 5-HAYED treated mouse. As shown by Tab.1, although AD brought about two folds ALT and AST increase to the mouse, 5-HAYED did not changed the odds, nor exerted the significant influences to sCR and BUN, because the indexes of sCR and BUN in the 5-HAYED-treated mouse were nearly the same as the control (Tab. 1).

**Tab. 1.** Clinical Chemistry and Blood Index Test. **Note:** AST: Aspartate Aminotransferase, ALT: Glutamic-Pyruvic Transaminase, RRs: Reference Ranges; SCr: Serum Creatinine, BUN: Blood Urea Nitrogen; RBC: Red blood cell count, MCV: Mean corpuscular volume, RDW: Red cell distribution width, HCT: Hematocrit, MCH: Mean corpuscular hemoglobin, MCHC: Mean corpuscular hemoglobin concentration, WBC: White blood cell count, NE: Neutrophil percentage, EO: Eosinophil percentage, BA: Basophile percentage, LY: Lymphocyte percentage, MO: Monocyte percentage, PLT: Platelet count, MPV: Mean platelet volume. ***N=6, p<5%***.

| Organ | Index | NC | 5-HAYED | AD | AD+5-HAYED | RRs [32] |
| --- | --- | --- | --- | --- | --- | --- |
| Liver | ALT (IU/L) | 80±7 | 78±6 | 167±8 ↑ | 174±13 ↑ | 54 ~242 |
|  | AST (IU/L) | 45±5 | 49±5 | 83±8 ↑ | 89±5 ↑ | 26 ~70 |
| Kidney | sCr (mg/dl) | 0.47± 0.02 | 0.49± 0.02 | 0.62± 0.02 | 0.52±0.04 | 0.30 ~1.00 |
|  | BUN (mg/dl) | 17.2±2.51 | 17.4±2.51 | 16.81±0.92 | 18.04±1.64 | 13.9 ~28.3 |
| Blood | RBC (10<sup>6</sup> count/μl) | 6.5±0.6 | 7.1±0.6 ↑ | 4.2±0.7 ↓ | 8.2±0.6 ↑ |  |
|  | MCV (fL) | 46.3 ±2.7 | 46.5 ±2.7 | 47.2±7.7 | 47.2±7.6 |  |
|  | RDW (%CV) | 24.1±2.9 | 24.5±2.9 | 25.2±4.2 | 26.1±5.4 ↑ |  |
|  | HCT (%) | 31.2 ±1.9 | 36.2 ±1.9 ↑ | 22.3±5.3 ↓ | 42.7±5.7 ↑ |  |
|  | MCHC (g/L) | 33.3 ±1.6 | 33.7 ±1.6 | 30.6±3.6 ↓ | 34.8±2.3 |  |
|  | WBC (10<sup>3</sup> count/μl) | 8.9±1.6 | 9.0±1.3 | 9.1±1. 5 ↑ | 9.3±2.1 ↑ |  |
|  | NE (%) | 18.0 ±3.0 | 18.1 ±3.0 | 18.2 ±4.2 | 17.8 ±4.5 |  |
|  | EO (%) | 2.6 ±0.3 | 2.5 ±0.4 | 2.8±0.6 ↑ | 2.4±0.7 ↓ |  |
|  | BA (%) | 2.1 ±0.4 | 2.2 ±0.3 | 2.3±0.4 | 2.3±0.6 ↑ |  |
|  | LY (%) | 83.3 ±3.6 | 84.8 ±2.7 | 96.6±2. 2 ↑ | 89.8 ±0.6 ↑ |  |
|  | MO (%) | 3.62 ±0. 35 | 3.87 ±0. 54 | 7.63±0.6 ↑ | 7.67±0.51 ↑ |  |
|  | PLT (10<sup>3</sup> count/μl) | 750±83.0 | 742±90.0 | 571±61 ↓ | 720±52 ↓ |  |
|  | MPV (fL) | 6. 7 ±0.7 | 6. 6 ±1.1 | 7.5 ±1.2 ↑ | 7.4±0.7 ↑ |  |

The AD attack and the introduction of 5-HAYED did not influence the mean corpuscular volume (MCV) or the percentage of neutrophils (NE). Although 5-HAYED did not ameliorate the AD-caused increases in red cell distribution width (RDW), monocyte percentage (MO) and white blood cell count (WBC), the indexes, including red blood cell count (RBC), hematocrit (HCT), eosinophil percentage (EO), basophile percentage (BA), platelet count (PLT), and mean corpuscular hemoglobin concentration (MCHC), which were decreased in the AD suffers, were recovered, and the high level of lymphocyte percentage (LY) and the mean platelet volume (MPV) in the AD mice were reversed after the 5-HAYED administration (Tab. 1), suggesting that 5-HAYED has an anti-inflammatory effect and can alleviate AD associated anemia to a extent.

## 3. Discussion

Alzheimer’s disease has a profound impact on patients, as well as their families and friends. However, the pathogeny is not yet fully understood. Gene mutations on PS1, PS2, APP and APOE have been associated with inherited Alzheimer’s disease [18, 19, 20]. In addition, environmental or lifestyle factors, such as heart disease, high blood pressure, and diabetes, may also play roles in the development of Alzheimer’s disease [21, 22, 23].

Free radicals corrupt the cell membrane, vessel wall, proteins, lipids or even DNA in cell nuclei [24]. In brain, iron is a fundamental catalyzer for free radical generation. The bivalent form of iron Fe^2+^ is capable of transferring one electron to O_2_, producing the superoxide radical •O_2_^-^. The reaction of Fe^2+^ with H_2_O_2_ produces the highly reactive hydroxyl radical (•OH). These oxygen free radicals, plus H_2_O_2_ and singlet oxygen, may imperil neurons in iron-rich regions [25]. Therefore, iron can be regarded as an important pathophysiologic element for AD.

The distribution and homeostasis of iron in brain may be regulated by transferrin, lactoferrin, iron responsive protein, ceruloplasmin, and ferritin. Among these regulators, transferrin is homogenously distributed around senile plaques and also found in astrocytes in the cerebral cortex of Alzheimer’s brain tissue. Most ferritin-containing cells are microglia associated with blood vessels in Alzheimer’s brain [26, 27]. Further, the permeability of veins in the choroid plexus hypothalamus (the blood brain barrier, BBB) also contains iron [28]. There, which are rich in iron-carrying proteins, iron is rich and radical is abundant and the oxidation of protein, nuclear DNA and lipids in neurons increase is understandable [29, 30, 31].

An agent capable of clearing radicals and redundant iron might be a potential AD treatment. Scavenging iron with DFO, DFP, M30 (5-[N-methyl-N-propargyaminomethyl]-8-hydroxyquinoline) or HLA20 (5-[4-propargylpiperazin-1-ylmethyl]-8-hydroxyquinoline) reportedly relives the symptoms of AD patients [32]. A NMDA (neuronal N-methyl-D-aspartate)–receptor antagonist was reported suppressed superoxide and peroxynitrite radicals in AD brains [33]. However, for the aspects of biochemistry, their effectiveness was severely limited. For example, DFO can tightly bind iron (III), but it failed to impede the progression of AD in clinical trials [34, 35]. DFP can cause infections by lowering neutrophils and white blood cell count [11, 12].

Aβ peptide is affinity to iron, it can bind iron ions at the sites D_1_, E_3_, and H_6, 13, 14_ [14], which chelating the excessive iron ions and lower the catalyzed free radicals in CSF. But for the scarce of iron-pro residues and low solubility, Aβ peptide is insufficient to purge the iron in AD brain. Even, Aβ_1-42_ peptide was presumed to spontaneously generate peptidyl radicals or hydrogen peroxide, which injuring neurons [36, 37].

The 5-HAYED peptide combined 5 times metal-pro amino residues histidine, glutamic acid and aspartic acid, moreover, it is high soluble for containing the polar amino acid tyrosine and the smallest nonpolar amino acid alanine. It is more effective in the iron scavenging and radical removal. By using it, we protected the neurons in the brain of AD mice, which brought about the cognition status amelioration and reverse AD associated anemia and inflammation. And by transferring 5-HAYED to KM mice, AD incidence lowered. Therefore, 5-HAYED peptide is a delightful iron scavenger to ease AD.

## 4. Conclusion

5-HAYED peptide can protect brain and decrease the incidence of AD by reducing the iron-catalyzed radicals via precipitating redundant free ions.

## 5. Materials and methods

### Materials

5-HAYED amino acid oligomers and the oligonucleotide encoding 5-HAYED amino acid oligomer were synthesized by GL Biochem Ltd. (Shanghai, China); pCEP4 and psisCAT6a Plasmids were constructed by Research Science Biotech Corp. (Shanghai, China); The SH-SY5Y cell line was obtained from American Type Culture Collection (ATCC, Manassas, VA, USA).

### Brain iron content topography scanning and measurement

The mouse brains were frozen coronal sectioned (100 μm thick) as reference [38]-figure.30 indexed. After weighing, the sections were placed on polycarbonate membranes and scanned by an X-fluorescence work station (Institute of High Energy Physics of China) for iron-topography.

The whole iron levels in brain, cell and medium were measured by an ICP (inductively coupled plasma emission spectrometer, Perkin Elmer Elan600, Fremont, CA, USA). Before it, they were individually freeze-dried, weighed and the nitrified. Gradient concentration FeCl_3_ solutions were used as standard samples. The experimental animals were treated under the Guidance for the Care and Use of Laboratory Animals of the National Institutes of Health and the protocol was approved by the Committee on the Ethics of Animal Experiments of Taizhou University (Permit Number: 13-1368).

### 5-HAYED oligomers observation by TEM

The synthesized 5-HAYED powder was dissolved in distilled water (4 mg/ml) and divided to two shares. The first was used as a control, without iron adding in. In the second, 5 μl 0.05 M FeCl_3_·6H_2_O solution was added into 20 μl 5-HAYED lyosol and then undergone a 24-hour incubation at 37°C. Then each sample was placed on a carbon film-coated copper mesh and with 2 min of 1% (m/v) phosphotungstic acid staining. After an air-dried, the samples were observed under a transmission electron microscope (TEM, H7650, Hitachi, Kyoto).

### Isothermal titration calorimetry (ITC)

To confirm the affinity between the iron atoms and 5-HAYED, purified 5-HAYED oligomers were exhaustively dialyzed against the ITC buffer. Following the dialysis, 5-HAYED oligomers were diluted to 200 μM, and FeCl_3_·6H_2_O was diluted to 1000 μM. The binding between iron and the 5-HAYED oligomers was measured using a VP-ITC Microcalorimeter (MicroCal, GE Healthcare). The heat of the ligand dilution was subtracted from the heat of the binding to generate a binding isotherm that was fit with a one-site binding model using Origin, allowing for the association constant (Ka), enthalpy (ΔH), and entropy (ΔS) of the interaction to be determined. Ka was used to calculate Kd (Kd=1/Ka).

### Infrared spectrum analysis

To determine how 5-HAYED binds iron ions, dried 5-HAYED-FeCl_3_ reactant was blended with K_2_Br powder; the IR spectra of the samples were measured by an infrared spectrometer (NEXUS870, NICOLET, USA). Pure 5-HAYED and K_2_Br blender were used as the controls.

### Cell culture

SH-SY5Y cells were cultured for determining how 5-HAYED protects cells from iron-induced cytotoxicity. They were cultured in RMPI DMEM medium at 37°C under 5% CO_2_ for 12 hours. And then divided to iron-stress (0.01 M FeCl_3_·6H_2_O) or iron-5-HAYED (containing 0.01 M FeCl_3_·6H_2_O and 100 pM 5-HAYED in the medium) group. The iron content is imitated the iron concentration in AD brain SCF, and the 5-HAYED content was optimized by previous tests. The with/without 5-HAYED (100 pM) group were set as the control. All plates of cells were cultured for 12 hours. Then, the cells in each dish were partially fixed with 2.5% glutaraldehyde for SEM; the remnants were submitted to flow cytometry for an apoptosis assay (BD FACSCalibur, Franklin Lake, USA).

### Cell apoptosis assay

Cells in each dish were collected individually and stained with Annexin-V-FITC and propidium iodide (PI) for 10 min in the dark at room temperature. A FACScan flow cytometer (Becton-Dickinson and company) was used to analyze cell apoptosis. The results were calculated using CellQuest Pro software (Becton, Dickinson and Company) to obtain the percentage of apoptotic cells from the total cells.

### Hydroxyl radical measurement

The whole brain of each mouse was ground on ice. Then the thoroughly ground samples were centrifuged at 10000 rpm for 15 min. The supernatants, along with culture medium (200 μl), were separately placed in 50 μl 1% salicylic acid solution (m/v). After the samples were incubated on a shaker at 37°C for 15 min, they were submitted to an enzyme-labeled meter (SpectraMax M5, Molecular Devices, USA) for transmittance value measurement at 510 nm. The OD value was set to index the hydroxyl radical level in each sample.

### Anti 5-HAYED immune serum preparing

Before the immunohistochemical test, 100 μl of 5-HAYED-physiological saline (0.5mg/ml) was mixed with 100μl incomplete Freund adjuvant, and then subcutaneous or intraperitoneal injected to each mouse every week. After two months, the whole blood of each mouse was extracted and the serum was isolated and frozen under −20°C for the later utility.

### 5-HAYED transgenic mouse building

5-HAYED amino acid oligomer coding DNA sequence (list in Fig.4-G) was constructed into pCEP4 plasmid. The PDGF promoter was combined upstream to initiate the 5-HAYED transcription. Then the plasmids (2μg/ml) were microinjected to the male prokaryotic of fertilized eggs derived from the donor mouse caged with the male. The well-growing eggs were in-transplanted to the fallopian tube of the pseudopregnant mouse. For the first 5-HAYED positive generation, the male and female mice were caged to mate reciprocally. The homozygotes of the second generation were used to statistic the AD incidence.

### PCR

To confirm the 5-HAYED transgenic mouse, the whole DNA was extracted from the mouse tail. Wherein, the PCR primers used for amplifying the 5-HAYED oligonucleotide are: the Forward: ATG CAT GCC TAC CAG GAT; and the Backward: ACT CTG GTA GGC ATG ATC.

PCR was performed in triplicate using a TaqMan PCR kit (TaqMan: CAS: N8080228) on an Applied Biosystems 7500 Sequence Detection System (Applied Biosystems, Foster City, CA, USA) under the following conditions: 10 min at 95°C, followed by 42 cycles of 15 sec at 95°C and 1 min at 60°C.

### Southern blot

The PCR products were electrophoresed on an agarose gel for separating by size. The DNA fragments in the gel were then transferred to a sheet of nitrocellulose membrane and the membrane was then exposed to a 5-HAYED hybridization probe (synthesized by Genepharm Ltd., Shanghai, China). After that, the membrane was washed for the excess probes removal and the pattern of hybridization is visualized on X-ray film.

### 5-HAYED immunohistochemical staining

Frozen brain slices were fixed for with 4% formaldehyde (v/v). After 3 washes with 0.1 M PBS, the sections were blocked with 0.1% BSA/PBS (m/v). After 1 hour, an anti-5-HAYED serum (diluted 1:1000 in PBS) was added to the sections. Incubated at 4°C overnight, the slices were washed and then covered with HRP labeled goat-anti-mouse IgG (diluted 1:300 in PBS). After 2 hours, the samples were washed again, and the substrate was bound by the enzyme DAB. After 30 min, the sections on the slides were stained with hematin. Washed for 3 times with PBS again, the samples were observed under a microscope. The sections contained scattered brown spots were 5-HAYED positive, indicating the introduction of 5-HAYED transgene.

### TUNEL assay

High iron-induced cell necrosis in brain tissue was assessed using a TUNEL kit (Order NO. E607172, Sangon Biotech Inc. Shanghai, China). In brief, the mouse brain was frozen sectioned (10 μm thickness), and then the slices were placed on slides. After incubating in Na-HEPES solution for 1 hour, they were treated with 10 mM H_2_O_2_ and 20 mM progesterone. Then they were washed in PBS and then permeabilized with 0.1% Triton-X100. Incubated in dark at 37°C for 1 hour in TUNEL reaction mixture, the samples were observed with microscopy. The bright green spots indicate the necrotic cells in the tissue.

### Morris water maze and functional magnetic resonance imaging (fMRI) assays

Morris water maze and fMRI assays were performed to evaluate the cognitive status of the tested mice. Specifically, the KM mice were divided into normal, wild aged AD model and 5-HAYED-treated groups. For the 5-HAYED-treated groups, 1.5 μM 5-HAYED oligomer contained saline was injected into the SCF weekly using stereotaxic coordinates of PA-1.0 mm, lateral-1.5 mm from bregma, and ventral-2.0 mm relative to dura. After 1 month, all groups of mice underwent a 4-day Morris water maze assay. The time spent and the distance swum to return the underwater platform was recorded to determine the individual cognitive status. Brain fNMRi was performed with an NMR spectrometer (E40, Flir Inc., USA) at 1.5 T for 30 min after the mouse was anaesthetized. The oxygen metabolism level in the brain was determined by the bright area in the encephalica.

### Hematological analyses

To evaluate the side effects of HAYED (5), mouse blood was collected according to the IFCC/C-RIDL protocol [31]. For the complete blood count, 1 ml of venous blood was drawn into a vacuum tube containing potassium 2 ethylene-diamine-tetraacetic acid (K_2_ EDTA). Hematological analyses were performed to evaluate the white blood cell count (WBC), neutrophil percentage (NE%), lymphocyte percentage (LY%), monocyte percentage (MO%), basophil percentage (BA%), eosinophil percentage (EO%), red blood cell count (RBC), hematocrit (HCT), mean corpuscular volume (MCV), mean corpuscular hemoglobin concentration (MCHC), red cell distribution width (RDW), platelet count (PLT) and mean platelet volume (MPV).

### Clinical chemistry

Clinical chemistry was performed to determine the side effects of 5-HAYED on the kidney or liver. Concretely, blood urea nitrogen (BUN) was assayed using a SpectraMax microtiter plate reader (Molecular Devices, LLC) and a MaxDiscovery Blood Urea Nitrogen Enzymatic Kit (Bioo Scientific Corporation). The Quantichrom Creatinine Assay Kit (BioAssay Systems) was utilized to measure creatinine in the serum (sCr). The serum activity of AST and ALT were determined using an automated analyzer (Selectra Junior Spinlab 100, Vital Scientific, Dieren, Netherlands) according to the manufacturer’s instructions.

### Data processing

Data are presented as the means ± SEM of three or more independent experiments, and differences were considered statistically significant at *p*< 0.05 using *t*-tests. The topographies of the iron distribution in the brains were processed with MATLAB 7.0 software (MathWorks, Natick, MA, USA).

## Acknowledgements

This study was supported by the Public Welfare Technology Research Grant for Zhejiang Social Development [2015C33248], Natural Science Foundation of Zhejiang Province [Y17H160027], Open object of the Key Laboratory of Shanghai Forensic Medicine [KF1606], Taizhou Science and Technology Program [1501KY32], Taizhou Science and Technology Program [1501KY32], Taizhou University Talent Fostering Fund [2015PY028].

## Appendix: Authors

| Name | Location | Role | Contribution |
| --- | --- | --- | --- |
| Zhenyou Zou | Taizhou University;<br>Purdue University | Author | Design and conceptualized study; analyzed the data; drafted the manuscript for intellectual content |
| Shengxi Shao | Imperial College London | Author | Carry out the IR, CD and ITC assays |
| Li Li | Netherland Cancer Institute | Author | Build up the HAYED transgenic mouse |
| Xiaoying Cheng | Taizhou University | Author | Carry out the Tunnel assay; radical and iron content measurements |
| Qiqiong Shen | Taizhou University | Author | Carry out the fMRI and MWM assay |

## Abbreviation

AD: Alzheimer’s disease
Aβ: Amyloid beta
KM mouse: Kunming mouse
NM: Normal
fNMRi: functional nuclear magnetic resonance

